# ONTdeCIPHER: An amplicon-based nanopore sequencing pipeline for tracking pathogen variants

**DOI:** 10.1101/2021.10.13.464242

**Authors:** Emira Cherif, Fatou Seck Thiam, Mohammad Salma, Georgina Rivera-Ingraham, Fabienne Justy, Theo Deremarque, Damien Breugnot, Jean-Claude Doudou, Rodolphe Elie Gozlan, Marine Combe

**Affiliations:** ISEM, Univ Montpellier, CNRS, IRD, Montpellier, France; Institut de Génétique Moléculaire de Montpellier, Université de Montpellier, CNRS, Montpellier, France; Université de Paris, Laboratory of Excellence GR-Ex, Paris, France; IRD, Centre IRD de Cayenne, Guyane Française

## Abstract

**Motivation:** Amplicon-based nanopore sequencing is increasingly used for molecular surveillance during epidemics (e.g. ZIKA, EBOLA) or pandemics (e.g. SARS-CoV-2). However, there is still a lack of versatile and easy-to-use tools that allow users with minimal bioinformatics skills to perform the main steps of downstream analysis, from quality testing to SNPs effect to phylogenetic analysis.

**Results:** Here, we present ONTdeCIPHER, an amplicon-based Oxford Nanopore Technology (ONT) sequencing pipeline to analyze the genetic diversity of SARS-CoV-2 and other pathogenes. Our pipeline integrates 13 bioinformatics tools. With a single command line and a simple configuration file, users can pre-process their data and obtain the sequencing statistics, reconstruct the consensus genome, identify variants and their effects for each viral isolate, infer lineage and, finally perform multi-sequence alignments and phylogenetic analyses.

**Availability and implementation:** ONTdeCIPHER is available at https://github.com/emiracherif/ONTdeCIPHER

**Supplementary information:** Supplementary data are available at …

## 1 Introduction

The SARS-CoV-2 pandemic resulted in over 231 million cases and 4.74 million deaths worldwide in less than two years (Dong et al., 2020). Since the first spark of the epidemic, global public health actors and international research teams have made considerable efforts that have resulted in unparalleled knowledge, from the identification of the viral genome to its biological mechanisms of infection and spread. A striking example is the first SARS-CoV-2 genome sequence published less than a month after the first viral infection report in December 2019, followed by thousands of genome sequences identified and published. Genome sequencing has enabled rapid identification of the virus, development of diagnostic tests and monitoring of the virus’ evolution (WHO, 2021). Nanopore sequencing technology combined with ARTIC’s Multiplex PCR-based protocol (Quick et al., 2017) is widely used to rapidly and reliably generate large amounts of raw sequencing data for molecular surveillance of the SARS-CoV-2 pandemic and the ZIKA, Ebola and Nipah outbreaks. However, there is still a lack of a versatile, practical tool that can perform the main steps of downstream analysis, from quality testing to SNPs effect to phylogenetic analysis, on a laptop or an HPC (high performance computing cluster) without the need for advanced bioinformatics skills, to meet the demand of recurrent analysis of hundreds of samples. In addition, few tools are specific to nanopore sequencing based on SARS-CoV-2 amplicons allowing genotyping and genome reconstruction (e.g. porCOV (Brandt et al., 2021), viralrecon (Patel et al., 2020) and interARTIC (Ferguson et al., 2021)). Here, we present ONTdeCIPHER, an amplicon-based Oxford Nanopore Technology (ONT) sequencing pipeline to analyze and assess the genetic diversity of SARS-CoV-2. ONTdeCIPHER integrates and orchestrates 13 bioinformatics tools combining ONT dedicated tools, SARS-CoV-2 specific tools and well-established generic tools. It is a stand-alone pipeline compatible with Ubuntu, mac and HPC distributions, including an easy-to-use configuration file. With a single command line and raw sequencing data as input, the users can pre-process their data and obtain sequencing quality, depth and coverage statistics. It then reconstructs consensus genomic sequences, identifies variants and their potential associated effects for each viral isolate, detects structural variants (SVs), infers lineage and finally performs multi-sequence alignments and phylogenetic analyses.

## 2 Features

While building on the main features of the ARTIC pipeline (https://github.com/artic-network/fieldbioinformatics), the ONTdeCIPHER pipeline incorporates additional useful features such as variant calling, variant annotation, lineage inference, multiple alignments and phylogenetic tree construction (Fig.1).

**Fig. 1.**
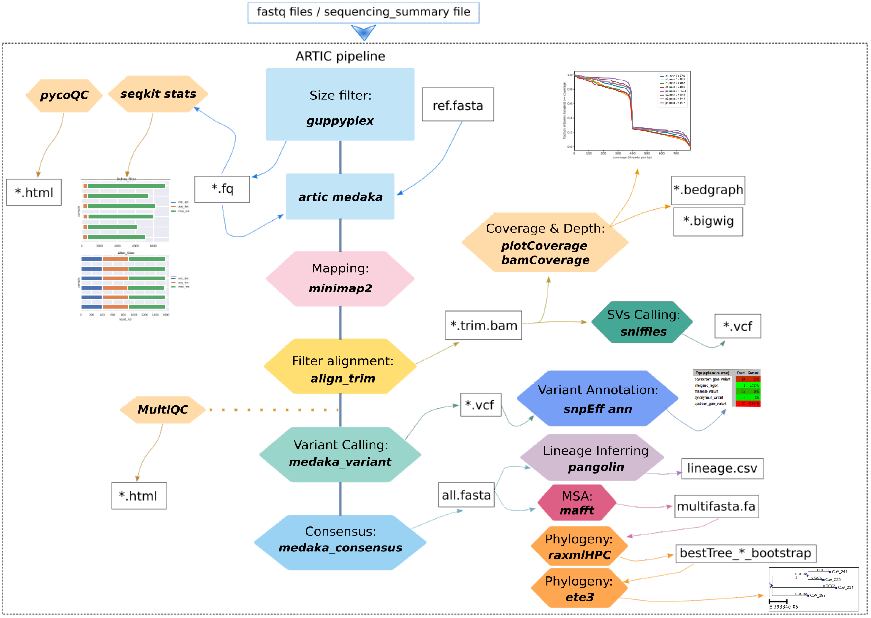
Overall ONTdeCIPHER pipeline. The diagram shows the ONTdeCIPHER features with the associated tool. All quality control features have the same color (dark yellow). Only the main results, files and figures are shown.

### Pre-processing and quality control

ONTdeCIPHER performs a pre-filtering step based on read length, followed by reads loss estimation using guppyplex *via* the ARTIC pipeline and SeqKit (Shen et al., 2016) respectively. The user can choose the maximum and minimum read length, with the default value of 400-700 bp (ARTIC primers V3). pycoQC is used by ONTdeCIPHER to calculate sequencing metrics (read length distribution, read quality scores, etc.). Then, using the ARTIC pipeline implemented in ONTdeCIPHER, the filtered reads are mapped to the Wuhan reference genome (Accession: NC_045512.2) using minimap2 (Li, 2018). Finally, amplicon dropouts during the multiplex PCR step and sequencing depth and coverage for each sample are estimated by ONTdeCIPHER using MultiQC (Ewels et al., 2016), plotCoverage (Ramírez et al., 2014) and bamCoverage (Ramírez et al., 2014), respectively.

### Genome reconstruction and genomic analysis

For each sample, ONTdeCIPHER reconstructs a reference-based consensus genome *via* the Medaka workflow and infers the viral lineage using the Pangolin Command-line tool (Rambaut et al., 2020). ONTdeCIPHER performs two variant calling steps for SNV (Simple Nucleotide Variation) and SV (Structural Variation) genotyping using Medaka and Sniffles (Sedlazeck et al., 2018). SNP variants are annotated using SnpEff (Cingolani et al., 2012). Finally, ONTdeCIPHER performs multiple alignments followed by phylogenetic tree construction for consensus genomic sequences obtained using the MAFFT command-line (Katoh et al., 2019), RAxML (Stamatakis, 2014) and ETE3 (Huerta-Cepas et al,, 2016).

## 3 Implementation

ONTdeCIPHER is an open-source pipeline coded in python3 compatible with Ubuntu distributions and macOS. It uses the Snakemake (Köster et al., 2012) features to perform reproducible and scalable data analysis (see Supplementary information Fig.S1) and seamlessly adapts to server, cluster, grid and cloud environments. ONTdeCIPHER’s dependencies and environments are installed and managed using the Conda package and environment management system. Currently, ONTdeCIPHER can be run locally or on an HPC.

ONTdeCIPHER includes 13 tools (see Supplementary information Table S1) validated by the bioinformatics community to perform key downstream analyses on raw sequencing data from amplicon-based ONT sequencing. In addition, it comes with a python master script *(run_deCiPHER.py)* allowing the user to run the various pipelines’ steps *via* a command line. ONTdeCIPHER requires two configuration files to run; 1) one containing the path to raw sequencing data (fastq and fast5 directories), the path to the sequencing_summary.txt file (optional), the minimum and maximum read lengths, the RaxML output name prefix (optional) and the version of the reference genome used, 2) the second is a tabulated file that contains the barcodes and the associated sample.

The main option of the master script is (--step). The user can select one of the following values: *pycoQC, pip_core, m_r_p* or *all. PycoQC* will calculate sequencing metrics and generate interactive QC plots, *pip_core* will run ONTdeCIPHER core pipeline (ARTIC pipeline, seqKit, DeepTools, Sniffles and snpEff), *m_r_p* will run mafft, raxml, ete3 and Pangolin while *all* will run all steps.

Output sub-directories are generated for each enabled step following a specific architecture (see the github repository for details). ONTdeCIPHER provides several visualization files (HTML, pdf, bedgraph, bigwig) to visualize and describe the results.

## 4 Applications

ONTdeCIPHER was tested on a subset of public clinical data of SARS-CoV-2 (Bioproject PRJNA675364) from Bull et al (2020) (see github repository test_data) and the SVs found mainly deletions, were comparable (see Supplementary information Table S2). ONTdeCIPHER was also tested on data from wastewater samples collected in French Guiana (South America) to assess the spatio-temporal dynamics and prevalence of the SARS-CoV-2 viral variants (see Supplementary information Fig. S2). ONTdeCIPHER was compared to the closest available tool, porCOV (Brandt et al., 2021) (see Supplementary information Table S3).

## 5 Conclusion and perspectives

ONTdeCIPHER uses a plethora of well-established tools to provide a practical pipeline for amplicon-based sequencing downstream analyses. ONTdeCIPHER aims to provide an efficient tool for large-scale monitoring of SARS-CoV-2 viral variants dynamics, but also of broader surveillance of a multitude of emerging environmental-based pathogens.

## Supporting information

see Supplementary information

## Acknowledgements

We acknowledge Dr. Stéphane Calmant, director of the IRD Centre at Cayenne, Dr. Mathieu Chouteau, researcher at CNRS Guyane, and Vincent Goujon, director of CNRS Guyane, for their logistical support and the availability of the laboratory facilities at Cayenne. We thank our local collaborators in French Guiana for their help to access wastewater samples (the Cayenne Prefecture, CTG, ARS, CACL, the town hall of Kourou, Sinnamary, Saint-Laurent, Apatou, Saint-Georges). We also acknowledge Marie-Ka Tilak for her precious advice, experience and discussions about viral RNA extractions and ONT library preparation and sequencing.

