## Supplementary material for "ONTdeCIPHER: An amplicon-based nanopore sequencing pipeline for tracking pathogen variants": see Supplementary information

+The authors wish it to be known that, in their opinion, the second and third authors should be regarded as Joint Second Authors.

**1 Snakemake Workflow**
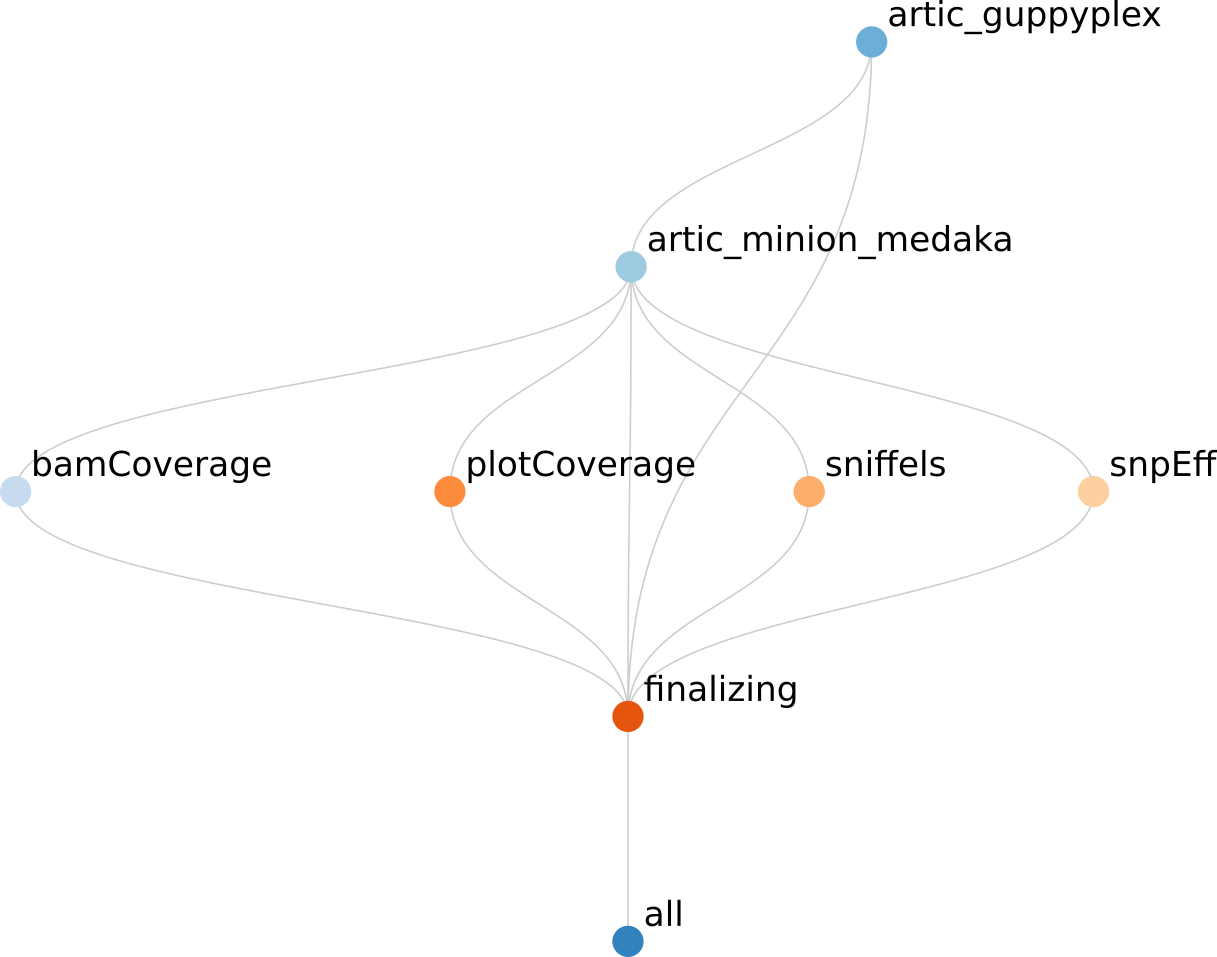

**Fig. S1.** ONTdeCIPHER snakemake workflow

**2 Implementation**

**Table S1** Tools implemented in ONTdeCIPHER

| Tools implemented in ONTdeCIPHER | Web site, Download site |
| --- | --- |
| **Quality control** | |
| seqKit | <https://bioinf.shenwei.me/seqkit/> |
| PycoQC | <https://tleonardi.github.io/pycoQC/)> |
| MultiQC | <https://multiqc.info/> |
| deepTools plotCovrage | <https://deeptools.readthedocs.io/en/develop/content/installation.html> |
| deepTools bamCovrage | <https://deeptools.readthedocs.io/en/develop/content/installation.html> |
| **Genome analysis** | |
| Artic* | <https://artic.network/ncov-2019/ncov2019-it-setup.html> |
| Minimap2 | <https://github.com/lh3/minimap2> |
| Sniffles | <https://github.com/fritzsedlazeck/Sniffles> |
| SnpEff | <http://pcingola.github.io/SnpEff/se_introduction/> |
| **MSA** | |
| MAFFT | <https://mafft.cbrc.jp/alignment/software/> |
| **Phylogeny** | |
| RAxML | <https://www.metagenomics.wiki/tools/phylogenetic-tree/construction/raxml> |
| ETE3 | <http://etetoolkit.org/> |
| **Lineage inference** | |
| Pangolin | <https://cov-lineages.org/pangolin.html> |

*****Are implemented in artic pipeline the ONT dedicated tools : Medaka, Nanopolish and guppyplex (Guppy)

**3 Examples of the results obtained with ONTdeCIPHER**

***Public data from Bull et al., 2020***

Four samples from Bull et al. (2020) for which SV results and raw data are available on Sequence Read Archive were analyzed with ONTdeCIPHER (all results are available in the OSF repository (<https://osf.io/jd2vz/?view_only=6d333ddc5a3045d297d5e3cc59e7e461>).

**Table S2** Structural variants detected by the *Step8_sniffles* of ONTdeCIPHER

| Specimen from Bull et al., 2020* | SVs detected with ONTdeCIPHER | Position_start | Position_end | Size | Supporting ONT reads |
| --- | --- | --- | --- | --- | --- |
| nCoV_214 | Deletion | 510 | 695 | 185 | 14 |
|  | Deletion | 23552 | 23583 | 31 | 59 |
| nCoV_249 | Deletion | 398 | 2455 | 2057 | 25 |
|  | Deletion | 1975 | 29352 | 27376 | 13 |
|  | Deletion | 2042 | 2970 | 928 | 24 |
|  | Deletion | 2087 | 4127 | 2040 | 1699 |
|  | Duplication | 2466 | 25560 | 23094 | 30 |
|  | Duplication | 2487 | 27701 | 25214 | 66 |
|  | Duplication | 2508 | 29172 | 26664 | 38 |
|  | Duplication | 2525 | 28650 | 26125 | 30 |
|  | Duplication | 4351 | 29861 | 25510 | 24 |
|  | Deletion | 4568 | 6611 | 2043 | 10 |
|  | Deletion | 8832 | 10821 | 1989 | 23 |
|  | Deletion | 10597 | 12357 | 1760 | 18 |
|  | Deletion | 11065 | 11276 | 211 | 33 |
|  | Deletion | 12815 | 15112 | 2297 | 20 |
|  | Deletion | 14855 | 16908 | 2053 | 18 |
|  | Deletion | 19250 | 21170 | 1920 | 12 |
|  | Deletion | 23495 | 25426 | 1931 | 30 |
|  | Deletion | 26780 | 26821 | 41 | 36 |
|  | Deletion | 27796 | 29173 | 1377 | 384 |
| nCoV_235 | Duplication | 2479 | 27556 | 25077 | 188 |
|  | Deletion | 7312 | 7515 | 203 | 12 |
|  | Deletion | 26779 | 26817 | 37 | 403 |
|  | Deletion | 27796 | 29173 | 1376 | 364 |
|  | Deletion | 29728 | 29777 | 49 | 113 |

*The nCov_183 was not presented in this table because no SVs were detected.

***Wastewater samples from French Guiana***
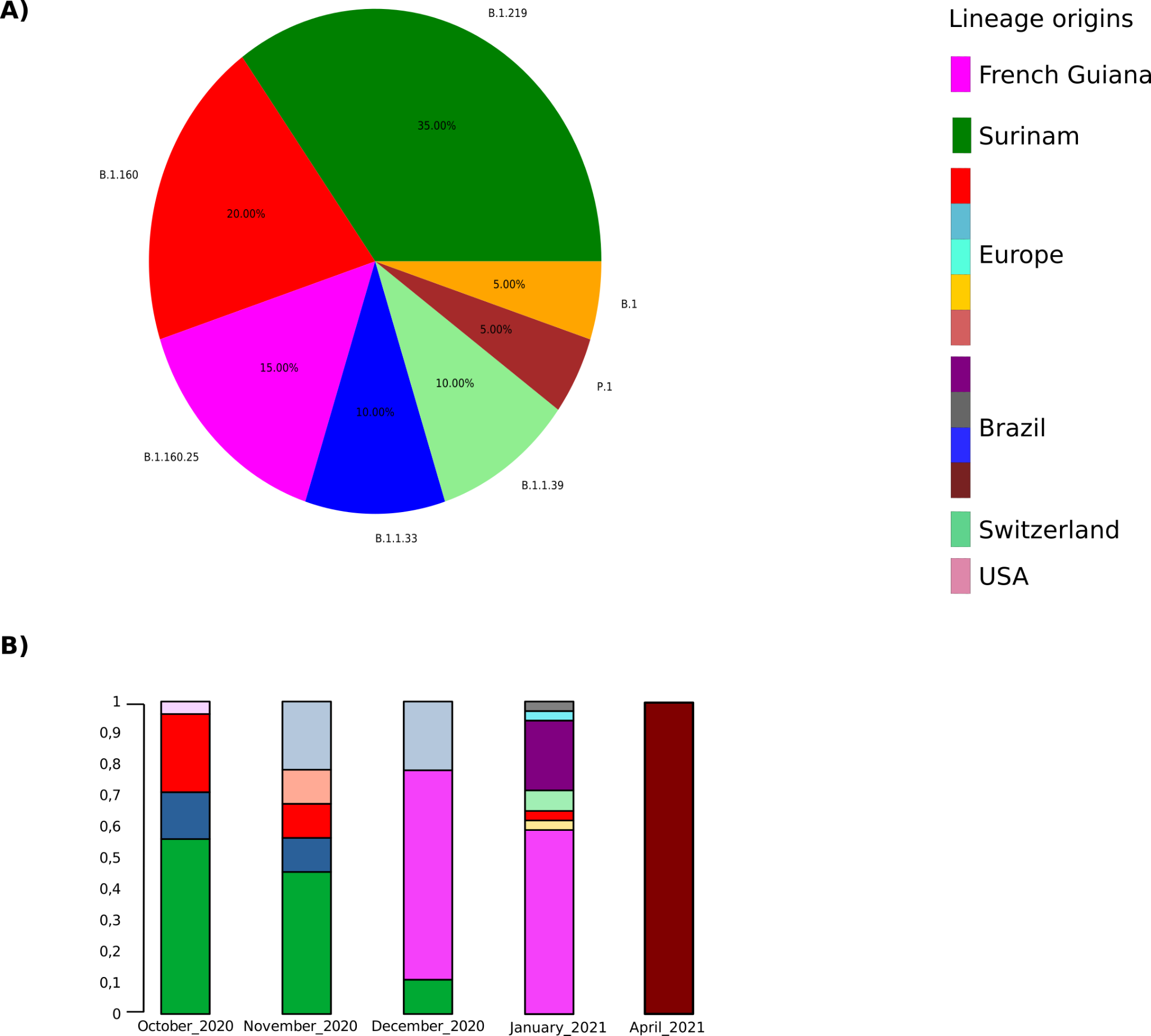

**Fig. S2** Spatio-temporal dynamics and prevalence of the SARS-CoV-2 variants in French Guiana

**3 Benchmark**

**Table S2** Comparision of the main features of ONTdeCIPHER and poreCov pipelines

|  | **ONTdeCIPHER** | **poreCov** |
| --- | --- | --- |
| **Analysis** | | |
| **Basecalling** | **✖** | **✓** |
| **Demultiplexing** | **✖** | **✓** |
| **Statistics on reads size** | **✓** | **✖** |
| **Sequencing coverage** | **✓** | **✖** |
| **Filtering by size** | **✓** | **✓** |
| **Quality information (QC)** | **✓** | **✓** |
| **Mapping** | **✓** | **✓** |
| **Variant Calling (SNVs)** | **✓** | **✓** |
| **Variant Calling (SVs)** | **✓** | **✖** |
| **Assembly** | **✓** | **✓** |
| **Variant annotation** | **✓** | **✖** |
| **Taxonomic classification** | **✖** | **✓** |
| **MSA & Phylogeny** | **✓** | **✓** |
| **Bootstrapping** | **✓** | **✖** |
| **SARS-CoV-2 lineage inference** | **✓** | **✓** |
| **Installation** | | |
| **OS** | **Linux, MacOS** | **Linux, MacOS** |
| **Workflow manager** | **Snakemake** | **Nextflow** |
| **Dependencies** | **Conda** | **Docker & Nextflow** |
| **Usage** | | |
| **Need Internet connection** | **No** | **Yes** |
| **Downloaded data** | **-** | **2.9 GB*** |
| **macOS Mojave (4 cores)+** | **12 min 43 sec** | **2h 30m 50s** |
| **Ubuntu 20.04 (10 cores)+** | **6 min 52 sec** | **39 min 12 sec** |

*Each time poreCov is launched the Kraken database is downloaded except if the user provides the path of the database.

+2.01Gb tested data.
